# High-throughput cancer hypothesis testing with an integrated PhysiCell-EMEWS workflow

**DOI:** 10.1101/196709

**Authors:** Jonathan Ozik, Nicholson Collier, Justin M. Wozniak, Charles Macal, Chase Cockrell, Samuel H. Friedman, Ahmadreza Ghaffarizadeh, Randy Heiland, Gary An, Paul Macklin

**Affiliations:** Argonne National Laboratory, Argonne, IL USA.; Dept. of Surgery, University of Chicago, Chicago, IL USA.; Opto-Knowledge Systems, Inc., Torrance, CA USA.; Lawrence J. Ellison Center for Transformative Medicine, University of Southern California, Los Angeles, CA USA.; Intelligent Systems Engineering, Indiana University, Bloomington, IN USA.

**Keywords:** Agent-based model, PhysiCell, cancer, immunotherapy, high throughput computing, EMEWS, hypothesis testing

## Abstract

**Background:** Cancer is a complex, multiscale dynamical system, with interactions between tumor cells and non-cancerous host systems. Therapies act on this combined cancer-host system, sometimes with unexpected results. Systematic investigation of mechanistic computational models can augment traditional laboratory and clinical studies, helping identify the factors driving a treatment’s success or failure. However, given the uncertainties regarding the underlying biology, these multiscale computational models can take many potential forms, in addition to encompassing high-dimensional parameter spaces. Therefore, the exploration of these models is computationally challenging. We propose that integrating two existing technologies—one to aid the construction of multiscale agent-based models, the other developed to enhance model exploration and optimization—can provide a computational means for high-throughput hypothesis testing, and eventually, optimization.

**Results:** In this paper, we introduce a high throughput computing (HTC) framework that integrates a mechanistic 3-D multicellular simulator (PhysiCell) with an extreme-scale model exploration platform (EMEWS) to investigate high-dimensional parameter spaces. We show early results in applying PhysiCell-EMEWS to 3-D cancer immunotherapy and show insights on therapeutic failure. We describe a generalized PhysiCell-EMEWS workflow for high-throughput cancer hypothesis testing, where hundreds or thousands of mechanistic simulations are compared against data-driven error metrics to perform hypothesis optimization.

**Conclusions:** While key notational and computational challenges remain, mechanistic agent-based models and high-throughput model exploration environments can be combined to systematically and rapidly explore key problems in cancer. These high-throughput computational experiments can improve our understanding of the underlying biology, drive future experiments, and ultimately inform clinical practice.

## Background

Cancer is a complex, dynamical system operating on many spatial and temporal scales: processes include molecular interactions (e.g., gene expression and protein synthesis; nanoseconds to minutes), cell-scale processes (e.g., cycle progression and motility; minutes to hours), tissue-scale processes (e.g., tissue mechanics and biotransport; minutes to days), and organ and organism-scale processes (e.g., organ failure and clinical progression; weeks, months, and years). Cancer-host interactions dominate throughout these scales, including interactions between tumor cells and the vasculature (hypoxic tumor cells trigger growth of new blood vessels; new but dysfunctional blood vessels supply further growth substrates and can promote metastasis), between tumor cells and stromal cells (tumor cells can prompt tissue remodeling that facilitates tissue invasion), and between tumor cells and the immune system (immune cells can kill tumor cells, but tumor cells can co-opt inflammation to promote their survival). See the reviews in [1, 2, 3, 4, 5, 6]. When designing and evaluating new cancer treatments, it is imperative to consider the impact on this complex multiscale cancer-host system.

Cancer-host interactions have been implicated in the poor (and sometimes surprising) clinical outcomes of existing and new treatments. Chemotherapies fail when molecular-scale processes (e.g., DNA repair failures, mutations, or epigenetic alterations) cause resistant tumor clones to emerge (multicellular-scale birth-death processes) which can survive the treatment [6, 7, 8, 9, 10, 11]. Anti-angiogenic therapies that target blood vessels were expected to be potent agents against cancer [12], but disrupting tissue perfusion inhibits drug delivery and increases hypoxia, which was subsequently shown to select for more aggressive tumor phenotypes, including alternative metabolism and increased tissue invasion [13, 14, 15]. On the other hand, medications originally developed for osteoporosis (bone loss) were found to reduce the incidence of bone metastases through unclear mechanisms, but hypothesized to arise from tumor-osteoclast interactions [16, 17, 18]. Such examples underscore the need to evaluate and improve cancer treatments from a cancer-host systems perspective.

Recent successes of cancer immunotherapies—such as CAR (chimeric antigen receptor) T-cell treatments [19, 20]—have brought heightened attention to cancer immunology. In some patients, immune cell therapies have been impressively successful, while other patient populations have demonstrated disappointing outcomes; this variability of patient response arises in part from the poorly-understood, complex interactions between cancer and the immune system [21, 22, 23, 24, 25, 26]. This suggests that better immune therapies could be designed through systematic investigations of tumor-immune interactions.

### Key elements for systematic and mechanistic investigation of cancer immunotherapy

Given the complexity and underlying uncertainty regarding the biological processes that drive cancer, dynamic computational models have been used to represent various cellular and molecular functions associated with cancer [27].

In particular, agent-based modeling [27] is an increasingly common computational modeling method that can aid in the translation of genetic/molecular/subcellular processes to the multicellular behavior of tumors and the host. Agent-based models (ABMs) can serve as modes for multiscale dynamic knowledge representation [28, 29], with the rules for each model representing a particular hypothesis of how the system may work. As such, they serve a potentially vital role in aggregating existing biological knowledge, and through simulation experiments exploring their behavior, can help establish the boundaries of the set of plausible hypotheses.

However, the dynamic multiscale models (e.g., ABMs) needed to approximate the complexity of the overall system are by their very nature resistant to formal analysis. Their overall behavior can only be evaluated by the execution of heuristic methods that require very large numbers of simulations, a process we term model exploration (ME). ME is a near-ubiquitous component in the development of models and algorithms; as applied to ABMs, it involves an iterative workflow where simulation experiments are carried out across a large range of parameter values (parameter space exploration) and varying perturbations and initial conditions (model behavior space exploration). Model outputs from a set of simulation experiments are evaluated against some predetermined metric, which informs the next iteration of simulation experiments. Advances in high-performance computing can allow the parallelization of this process, resulting in high-throughput dynamic knowledge representation and hypothesis evaluation to address a current bottleneck in the Scientific Cycle [30]. However, we propose that the ME process itself can be enhanced with a computational framework for its workflow [31].

In this paper, we formulate the requirements for a computational experimental system for systematic, high-throughput hypothesis testing and optimization. We provide an example of how high-throughput hypothesis testing can be applied to the complex problem of tumor-immune interactions using a novel framework that integrates a multiscale mechanistic model development platform—PhysiCell [32] and BioFVM [33]— within a computational ME manager—Extreme Model Exploration with Swift (EMEWS) [31].

We then present early work on implementing our proposed high-throughput hypothesis testing and optimization framework with PhysiCell and EMEWS. After an initial 2-D test deployment that explored the impact of tumor oxygenation, we present a high-throughput investigation of a 3-D computational model of the adaptive immune response to tumor cells from [32]. This work exposed new and counter-intuitive insights on tumor-immune cell attachment dynamics and the nonlinear role of immune cell homing on successful and unsuccessful tumor suppression. The study performed over 1.5 years’ worth of computational investigation in just over two days—a feat that is computationally infeasible without a framework that merges mechanistic modeling with efficient model exploration.

We close with a discussion of our ongoing and future work to implement the full PhysiCell-EMEWS framework *iterative* hypothesis exploration and optimization, along with potential applications in developing synthetic multicellular cancer treatment systems. We note that both PhysiCell and EMEWS are free and open source software. PhysiCell is available at http://PhysiCell.MathCancer.org and EMEWS is available at http://emews.org.

## Method

### 3-D cancer immunology model exploration using PhysiCell-EMEWS

There have been multiple projects utilizing agent-based/hybrid modeling of tumors and their local environments [34, 35, 36, 37]. Review of this work and our own has led to the following list of key elements needed to systematically investigate cancer-immune dynamics across high-dimensional parameter/hypothesis spaces to identify the factors driving immunotherapy failure or success:

1. efficient 3-D simulation of diffusive biotransport of multiple (5 or more) growth substrates and signaling factors on mm3-scale tissues, on a single compute node (attained via BioFVM [33]);
2. efficient simulation of 3-D multicellular systems (10^5^ or more cells) that account for basic biomechanics, single-cell processes, cell-cell interactions, and flexible cell-scale hypotheses, on a single compute node (attained via PhysiCell [32]);
3. a mechanistic model of an adaptive immune response to a 3-D heterogeneous tumor, on a single compute node (introduced in [32]);
4. efficient, high-throughput computing frameworks that can automate hundreds or thousands of simulations through high-dimensional hypothesis spaces to efficiently investigate the model behavior by distributing them across HPC/HTC resources (attained via EMEWS [31]); and
5. clear metrics to quantitatively compare simulation behaviors, allowing the formulation of a hypothesis optimization problem (see Proposition: hypothesis testing as an optimization problem).

### Efficient 3-D multi-substrate biotransport with BioFVM

In prior work [33] we developed BioFVM: an open source framework to simulate biological diffusion of multiple chemical substrates (a vector *ρ*) in 3-D, governed by the vector of partial differential equations (PDEs)

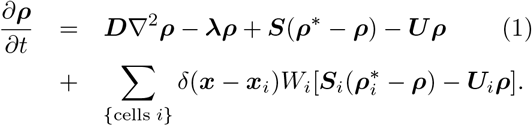

Here, **D** is the vector of diffusion coefficients, **λ** gives the decay rates, **S** and **U** are vectors of bulk source and uptake rates, and for each cell *i*, S_*i*_ and U_*i*_ are its secretion and uptake rates, *W_i_* is its volume, and x_*i*_ is its position. All vector-vector products (e.g., *λρ*) are component-wise, *ρ∗* denotes a vector of saturation densities (at which secretion or a source ceases), and *δ* is the Dirac delta function.

As detailed in [33], we solve this equation by a first-order operator splitting: we solve the bulk source and uptake equations first, followed by the cell-based sources and uptakes, followed by the diffusion-decay terms. We use first-order implicit time discretizations for numerically stable first-order accuracy. When solving the bulk source/decay term, we have an *independent* vector of linear ordinary differential equations (ODEs) in each computational voxel of the form:

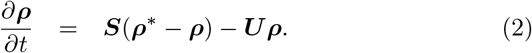

Each of these sets of ODEs can be solved with the standard backwards Euler difference, giving a firstorder accurate, stable solution. We trivially parallelize the solution by dividing the voxels across the processor cores with OpenMP: each thread works on a single voxel’s set of ODEs. Moreover, we wrote the ODE solver to work vectorially, with a small set of BLAS (basic linear algebra subprograms) implemented to reduce memory allocation, copy, and deallocation operations. (We implemented specific BLASes as needed to keep the framework source small and minimize dependencies to facilitate cross-platform portability across Windows, Linux, OSX, and other operating systems.) We solved the cell-centered sources and sinks similarly, by dividing the solvers across the cells by OpenMP (one set of ODEs per cell); note that each cell will act on the substrates in the voxel containing the cell center, by the Dirac delta formulation.

We solve the diffusion-decay equation by the locally one-dimensional (LOD) method, which transforms a single 3-D PDE into a series of three 1-D PDEs (one PDE with respect to the *x* derivatives, one for the *y* derivatives, and one for the *z* derivatives) [38, 39]. In any *x* -, *y* -, or *z*-strip, using centered 2^nd^-order finite differences for the spatial derivative and backward 1^st^-order Euler differences yields a tridiagonal linear system for each substrate’s PDE; because each PDE has the same form, we have a vector of tridiagonal linear systems. In [33], we solved this system with a vectorized Thomas algorithm [40]: an efficient O(*n*) direct linear solver for a single tridiagonal linear system, which we vectorized by performing all addition, multiplication, and division operations vectorially (with term-wise vector-vector multiplication and division). As a further optimization, we took advantage of that fact that **D** and ***λ*** are constant and noted that the forward sweep stage of the Thomas algorithm only depends upon **D**, ***λ***, and the spatial mesh, but not on the prior or current solution. Thus, we could pre-compute and cache in memory the forward-sweep steps in the *x-*, *y-*, and *z*-directions to reduce the processing time. We tested on numerous computational problems, and found the overall method was first-order accurate and stable in time, and second-order accurate in space [33]. Moreover, we found that the computational speed scaled linearly in the number of PDEs solved, with a slope much less than one: Simulating 10 PDEs takes approximately 2.6 times more computational effort than a single PDE, whereas sequentially solving 10 PDEs requires approximately 10 times more effort than a single PDE. See further results in [33].

In testing, we have found that this system can simulate 5-10 diffusing substrates on 1 million computational voxels (sufficient to simulate 8 mm3 at 20 *µ*m resolution) on a quad-core desktop workstation with 2 GB of memory; the performance was faster on a single compute node with greater computational core counts. This CPU-based algorithm maximizes cross-platform compatibility, but we anticipate a GPU implementation would be at least an order of magnitude faster.

### Efficient 3-D multicellular simulations with PhysiCell

In [32], we developed a 3-D agent-based modeling framework by extending BioFVM’s *basic agents* (discrete cell-like agents with static positions, which could secrete and consume chemical substrates in the BioFVM environment) to create extensible software cell agents. Each cell has an independent, hierarchically-organized phenotype (the cell’s behavioral state and parameters) [41, 42]; user-settable function pointers to define hypotheses on the cell’s phenotype, volume changes, cell cycling or death, mechanics, orientation, and motility; and user-customizable data. The cells’ function pointers can be changed at any time in the simulation, allowing dynamical cell behavior and even switching between cell types. The overall program flow progresses as follows. In each time step:

1. Update the chemical diffusing fields by solving the PDEs above with BioFVM.
2. For each cell, update the phenotype by evaluating each cell’s custom phenotype function. Also run the cells’ cell cycle/death models, and volume update models. This step is parallelized across all the cells by OpenMP.
3. Serially process the cached lists of cells that must divide, and cells that must be removed (due to death). Separating this from step 2 preserved memory coherence.
4. For each cell, evaluate the mechanics and motility functions to calculate the cells’ velocities. This step can be parallelized by OpenMP because the cell velocities are based upon relative positions.
5. For each cell, update the positions (using the second-order Adams-Bashforth discretization) using the pre-computed velocities. This step is also parallelized by OpenMP.
6. Update time.

The cell velocity functions (adapted from [35]) requires computing *n-1* pairwise cell-cell mechanical interactions for all *n* cells, giving O(*n*^2^) computational performance—this would be prohibitive beyond 103 or 10^4^ cells. However, biological cells have finite interaction distances, so we created an interaction testing data structure that placed each cell’s memory address in a Cartesian mesh, and limited cell-cell mechanical interaction testing to the nearest interaction voxels. This reduced the computational effort to *O* (*n*). PhysiCell uses separate time step sizes for biotransport (Δ*t ∼* 0.01 min), cell mechanics (Δ*t ∼* 0.1 min), and cell processes (Δ*t ∼* 6 min) to take advantage of the multiple time scales. See [32] for further details.

### Extreme-scale Model Exploration with Swift (EMEWS)

While detailed modeling approaches like PhysiCell allow higher fidelity representation of molecular, cellular, and tissue dynamics in cancer, they present substantial challenges. These challenges center on dealing with the large parameter spaces of these models and the highly nonlinear relationship between ABM input parameters and model outputs due to multiple feedback loops and emergent behaviors. Since their complexity limits the use of formal analytical approaches, the calibration and interpretation of complex ABMs often requires heuristic model exploration approaches that adaptively evaluate large numbers of simulations. These approaches often involve complex iterative workflows driven by sophisticated ME algorithms, such as genetic algorithms [43] or active learning [44, 45], which adaptively refine model parameters through the analysis of recently generated simulation results and launch new simulations.

The Extreme-scale Model Exploration with Swift (EMEWS) framework [31] is built on the the general-purpose parallel scripting language Swift/T [46], and is used to generate dynamic, highly concurrent simulation workflows for guiding ABM exploration in highdimensional parameter spaces. EMEWS enables the direct integration of external ME algorithms to control and coordinate the running and evaluation of large numbers of simulations via iterative HPC workflows. The general-purpose nature of the underlying Swift/T workflow engine allows the supplementing of the workflows with additional analysis and post-processing as well.

EMEWS enables the user to plug in both ME algorithms and scientific applications, such as PhysiCell ABMs. The ME algorithm can be expressed in Python or R, utilizing high-level queue-like interfaces with two implementations: EQ/Py and EQ/R (EMEWS Queues for Python and R). The scientific application can be implemented as an external application called through the shell, in-memory libraries accessed directly by Swift (for faster invocation), or Python, R, Julia, C, C++, Fortran, Tcl and JVM language applications. Thus, researchers in various fields who may not be parallel programming experts can simply apply existing ME algorithms to their existing scientific applications and run large-scale computational experiments without explicit parallel programming. A key feature of this approach is that neither the ME algorithm nor the scientific application is modified to fit the framework.

### Mechanistic 3-D model of adaptive immune response to heterogeneous tumors

#### Heterogeneous tumor

In [32], we developed an initial model of an adaptive immune response to a heterogeneous tumor. In the model, each cell exchanges cell-cell adhesive and “repulsive” forces, and enters the cell cycle at a rate that increases with oxygen availability. Each cell consumes oxygen, which diffuses from the simulation’s boundary voxels, leading to the formation of hypoxic gradients. Where oxygenation drops to very low levels, tumor cells become necrotic and slowly lose volume. To model heterogeneity, each cancer cell has a normally distributed mutant “oncoprotein” expression 0 *≤ p ≤* 2 (with mean 1, standard deviation 0.3). Cells with greater expression of *p* are modeled as entering the cell cycle more rapidly. See [32] for more details and references.

#### Immunogenicity and immune response

As a simplified model of MHC (major histocompatibility complex: a surface complex that presents a “signature” sampling of fragments of the cell’s peptides, allowing immune cells to learn to recognize the body’s own cells [47, 48]), we assume cells with greater *p* expression are more immunogenic: more likely to present abnormal peptides on MHC and be recognized as targets for immune attack. All tumor cells secrete an immunostimulatory factor that diffuses through the domain. (Even *in situ* tumors are known to prompt immune cell homing [49].) Immune cells perform biased random migration (chemotaxis) along gradients of this factor, test for collision with cells, and form tight adhesions with any cells that are found.

For any time interval [*t, t* + Δ*t*] while an immune cell *i* is attached to another cell *j*, the immune cell attempts to induce apoptosis (programmed cell death) with probability *ripj* Δ*t*, where *ri* is the immune cell’s killing rate for a normal immunogenicity (*p* = 1), and *pj* is the *j*th cell’s oncoprotein expression; this models activation of a death receptor, such as FAS. For more background biology and references, see [32]. If an immune cell triggers apoptosis, it detaches and continues its search for new immunogenic targets. Otherwise, it remains attached, but with a similar stochastic process to regulate how long it remains attached.

#### Sample 3-D simulation

In [32], we simulated this problem in 3D for an initial cell population of approximately 18,000 cells in a *∼* 5 mm3 domain on a quad-core desktop workstation. At the simulation start, tumor cells are very heterogeneously distributed; see the first frame in Figure 1, where the tumor cells are shaded by *p* expression from blue (*p ≤* 0.5) to yellow (*p ≥* 1.5). By two weeks (Figure 1, third frame), the tumor has grown by an order of magnitude (from *∼*104 to 105 cells), there is clear selection for the cells with the most *p* (the tumor is visibly more yellow), oxygen transport limits have lead to the formation of a necrotic core (brown central region), and the initial spherical symmetry has been lost due to the formation of clonal foci (larger, more homogeneous yellow regions).

**Figure 1.**
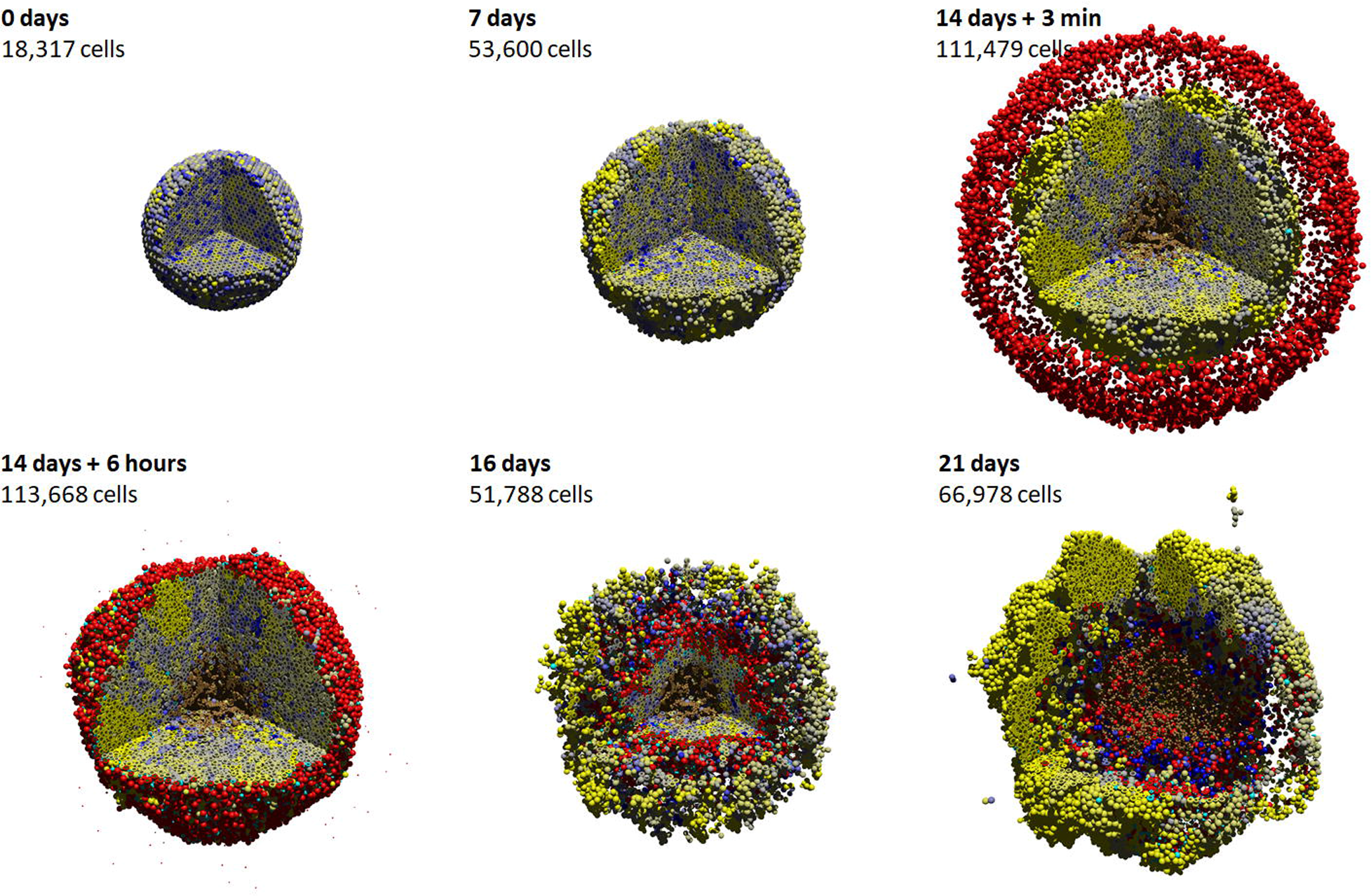
Sample 3-D cancer-immune simulation. 3-D simulation of adaptive immune response to a heterogeneous tumor (cells ranging from blue (low proliferation and immunogenicity) to yellow (high proliferation and immunogenicity). Immune cells are red; cyan cells have undergone apoptosis due to immune attack. A high-resolution animation can be viewed at https://www.youtube.com/watch?v=nJ2urSm4ilU. Adapted with permission (via CC-BY 4.0) from [32].

At this point, we introduced 7500 immune cells (red) and applied the immune response model. By later simulation times (16 and 21 days in Figure 1), we observed that the immune cells continue migrating along the chemical gradient until reaching the center where the gradient is approximately flat. Due to the particular choice of motility parameters for the immune cells, they become temporarily trapped in the center, allowing tumor cells to evade therapy and re-establish the tumor. A highresolution video of this simulation can be viewed at https://www.youtube.com/watch?v=nJ2urSm4ilU.

### Proposition: hypothesis testing as an optimization problem

We posit that the application of an integrated framework where the PhysiCell model is deployed within the EMEWS framework can be used to take advantage of EMEWS’s more advanced ME capabilities to inform hypothesis exploration as a function of parameter space search (e.g., via active learning) and hypothesis optimization (e.g., via genetic algorithms). As an example, we describe the following set of parameters that represent a space of possible interactions governing tumor-immune interactions, and how that space could be explored:

1. A family of cell behavior hypotheses and constraints on their parameter values. For example:

a. immune cells can exhibit any combination of random motility, chemotaxis towards tumor cells, or chemotaxis away from other immune cells
b. attached immune cells can secrete immunoinhibitory or immunostimulatory factors
c. tumor cells can secrete immunoinhibitory factors, but at a cost to cellular energy available for proliferation
d. the microenvironment can have variable far-field oxygenation values.
2. A mechanistic computational model for simulating the cancer-host system under the hypotheses. For example:

a. We implement the additional diffusion equations in BioFVM.
b. We implement the prior tumor cell immunogenicity model, and add a basic model of cell metabolism (e.g., as in [50]) with extra energy cost for secreting the immunoinhibitory factor.
c. We implement the prior immune cell adaptive response model but vary the cell motility according the specific hypotheses for migration bias along the various chemical gradients, the level of randomness, and we vary decrease the migration speed, adhesion rate, and cell killing rate under immunoinhibition.
3. A set of target system behaviors and/or validation data. For example:

a. We seek hypotheses that result in emergence of immune-resistant tumors.
4. A model error metric to compare models and assess their match to target behavior. For example:

a. For a set of hypotheses, we quantify the number of tumor cells after 48 hours of immune attack, the secretion level of the immunoinhibitory factor, and the mean immunogenicity (mutant oncoprotein).

Given these user inputs, the proposed PhysiCell-EMEWS system would distribute simulations across the hypothesis space (each running independently on its own compute node, where they are optimized). For succinctness, we refer to a point in the hypothesis space as a single *simulation ruleset*. Because these models are stochastic, EMEWS will initialize multiple simulations for each ruleset. EMEWS then collects the simulation outputs, evaluates the user-supplied metric against the target model behavior, and either reports the best hypothesis ruleset (if only one iteration is allowed), or repeats the process to refine the current best hypothesis ruleset (e.g., by a genetic algorithm). Each iteration is a high-throughput hypothesis test. And the overall iteration is hypothesis optimization. See Figure 2.

**Figure 2.**
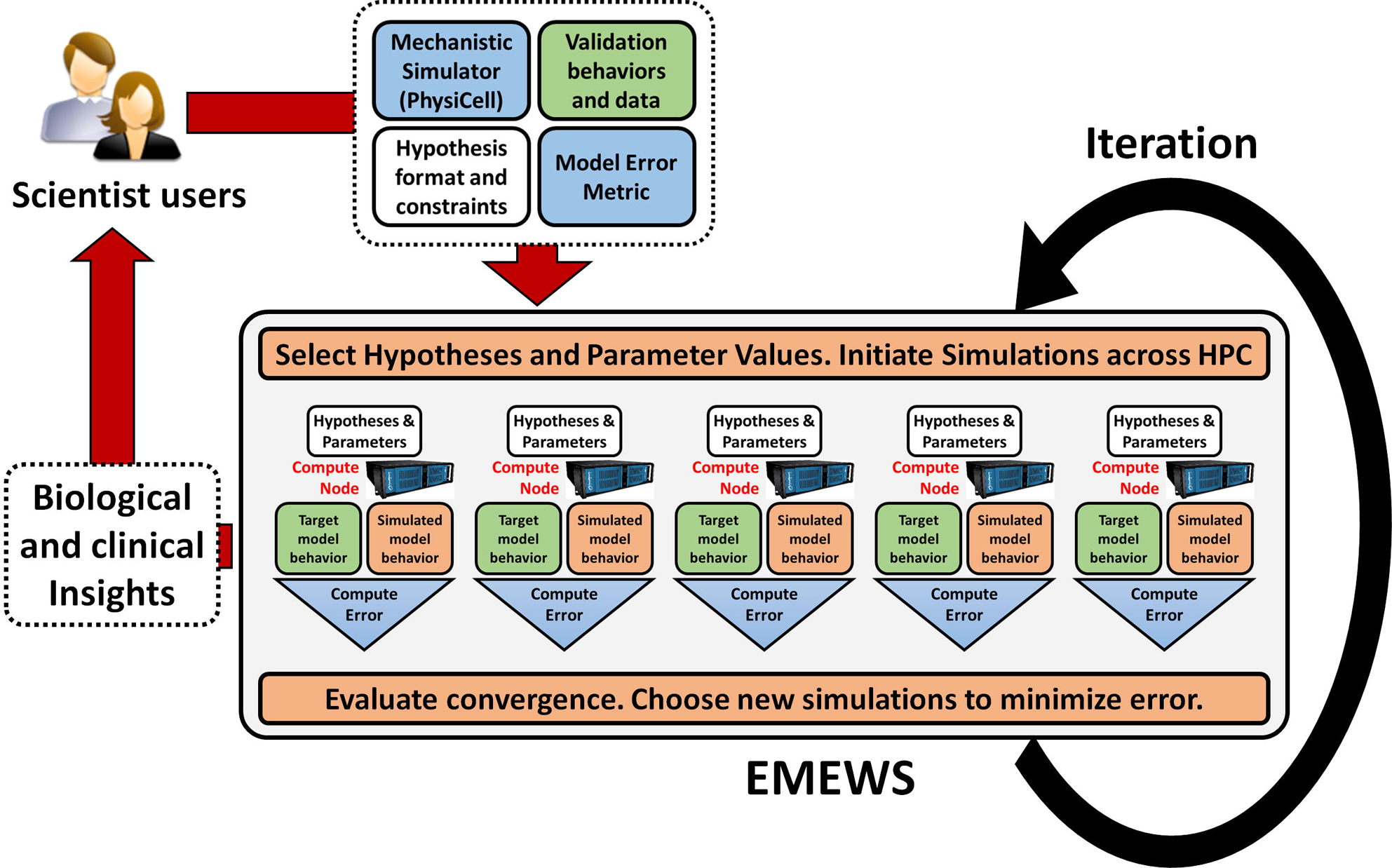
Hypothesis testing as an optimization problem. If scientific users can (1) formulate a range of hypotheses, (2) supply an efficient 3-D mechanistic simulator (BioFVM+PhysiCell), (3) provide validation behaviors and/or data, and (4) supply an error metric, then the combined PhysiCell-EMEWS system can automatically explore the space of hypotheses, initiate simulations on HPC/HTC resources, collect data to evaluate the error metric, and then make further decisions on which hypotheses and parameter values to explore next. The framework iteratively sharpens hypotheses that bring new biological and clinical insights.

The output is a set of hypotheses *H* that lead to the desired cell behaviors. For example, in hypoxic conditions, we may see less selection for the immunoinhibitory secreting cells due to limited nutrients, unless the cells are under attack by many immune cells. This hypothesis could then be tested experimentally. If the hypothesis does not hold experimentally, we would refine the computational model (e.g., focusing more on hypoxic cell metabolic and motile adaptations.)

## Results

We now demonstrate the first steps in implementing and testing the PhysiCell-EMEWS hypothesis optimization system: we conduct a single iteration of ME on a 2-D hypoxic cancer study, and then we test the 3D cancer-immune model on a high-throughput study that reduced over a year of continumous computing time to just 2 days.

### Test deployment of PhysiCell within EMEWS

The initial example of integrating PhysiCell with EMEWS involved examination of the effect of hypoxic conditions on tumor growth. This involved the development of a fast 2-D tumor simulator that could simulate 48 hours of oxygen-limited tumor growth in 1-2 minutes. The framework integration proceeded as in the Proposition: hypothesis testing as an optimization problem above. To work through user-supplied elements:

1. Oxygenation conditions could vary from completely anoxic (0 mmHg) to typical values of well-oxygenated breast tissue (60 mmHg; see [33, 51]). The initial cell population could vary from 1 to 2000 cells.
2. PhysiCell was used to create a program that could read these two hypothesis parameters at the command line, initialize the simulation, and run to 48 hours without user input.
3. The target behavior was to maximize live cell fraction.
4. The model metric was the live cell fraction after 48 hours.

We implemented a parameter sweep of PhysiCell using EMEWS, with the following oxygenation values:

0, 2.5, 5, 8, 10, 15, 38 or 60 mmHg

and the following initial cell counts:

1, 10, 100, 1000, 2000

EMEWS saved the model outputs in separate directories, facilitating subsequent postprocessing analysis and visualization. We plot a 2-D array of the final simulation images in Figure 3 and the final live cell counts in Figure 4 (top). As expected, increasing the initial cell count always increases the final cell count (and overall tumor size) 48 hours later, but for any fixed oxygenation condition, this also leads to greater prevalence of necrosis, and a nonmonotonic effect on final live cell fraction (Figure 4 (bottom)).

**Figure 3.**
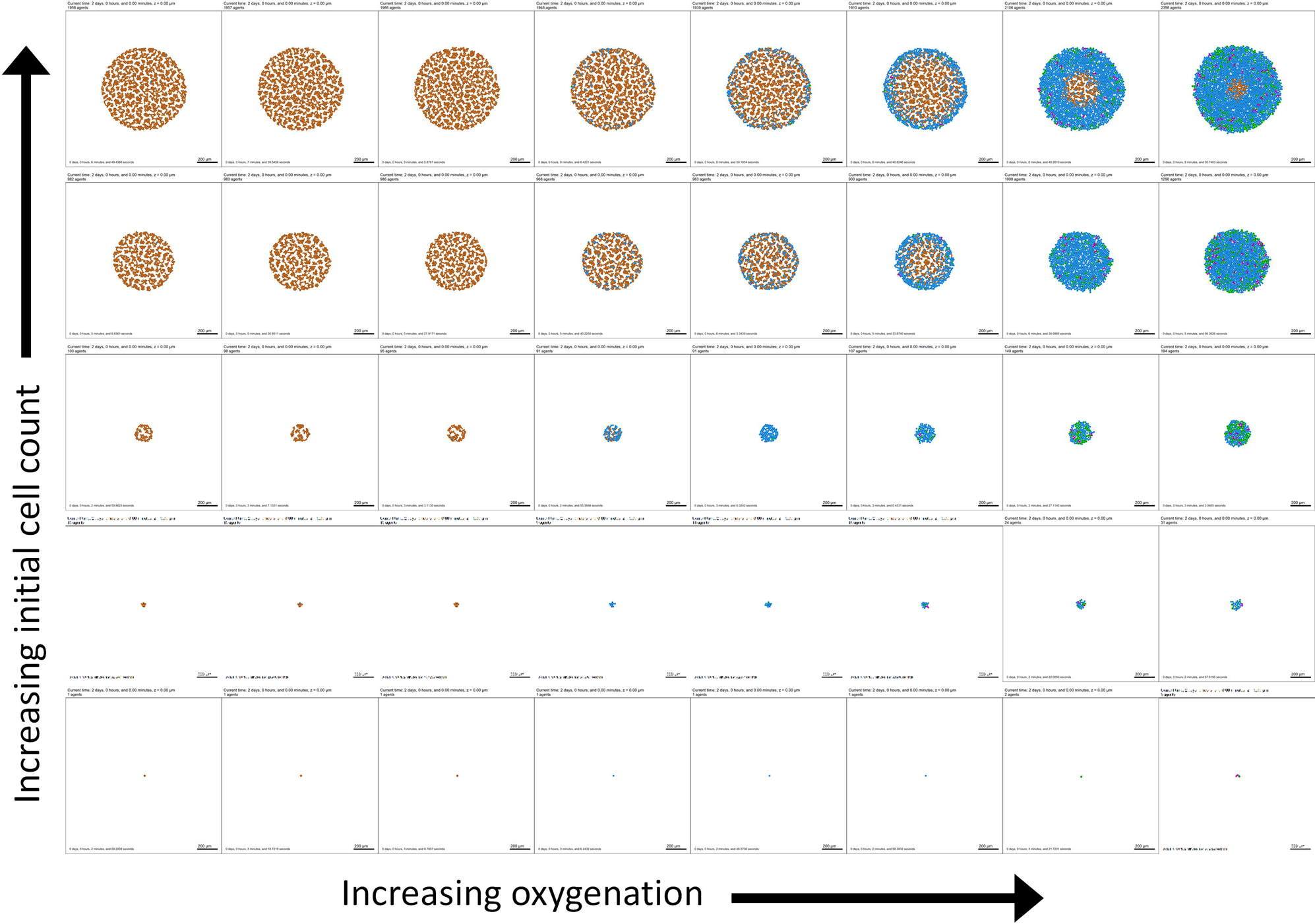
First PhysiCell-EMEWS test on cancer hypoxia: tumor plots. Here necrotic cells (dead by oxygen starvation) are brown, non-cycling cells are blue, and cycling cells are green and magenta. Increasing the initial cell count increases the final cell count, but also increases the final dead cell fraction (seen as the increasing prevalence of brown).

In Figure 4 (bottom), we plot the final live cell fraction as a function of the initial cell count, for each fixed oxygenation condition. For low oxygenation conditions (0, 2.5 mmHg), almost all cells are dead at 48 hours regardless of cell seeding choices. For intermediate oxygenation conditions (5 to 38 mmHg), the effect is nonmonotonic: for small initial cell populations (1 or 10 cells), stochastic apoptosis effects can sometimes leave a smaller final live fraction than a larger cell population; this highlights the importance of testing multiple simulation replicates for stochastic models. Past 100 initial cells, the stochastic effects are reduced, and increasing the initial cell count results in a lower final live fraction (due to oxygen depletion by the larger cell population and the emergence of a necrotic core). In particular, for these simulations increasing from 1000 to 2000 cells decreased the final live cell fraction. This behavior was not observed for high oxygenation (60 mmHg): no portions of the tumor ever drop below the necrotic threshold. Moreover, this simulated cell line has saturating proliferation above 38 mmHg pO2 (tissue physioxia [51] and so for sufficiently high initial oxygenation, the entire tumor stays about this threshold where there is no oxygen constraint to growth.

**Figure 4.**
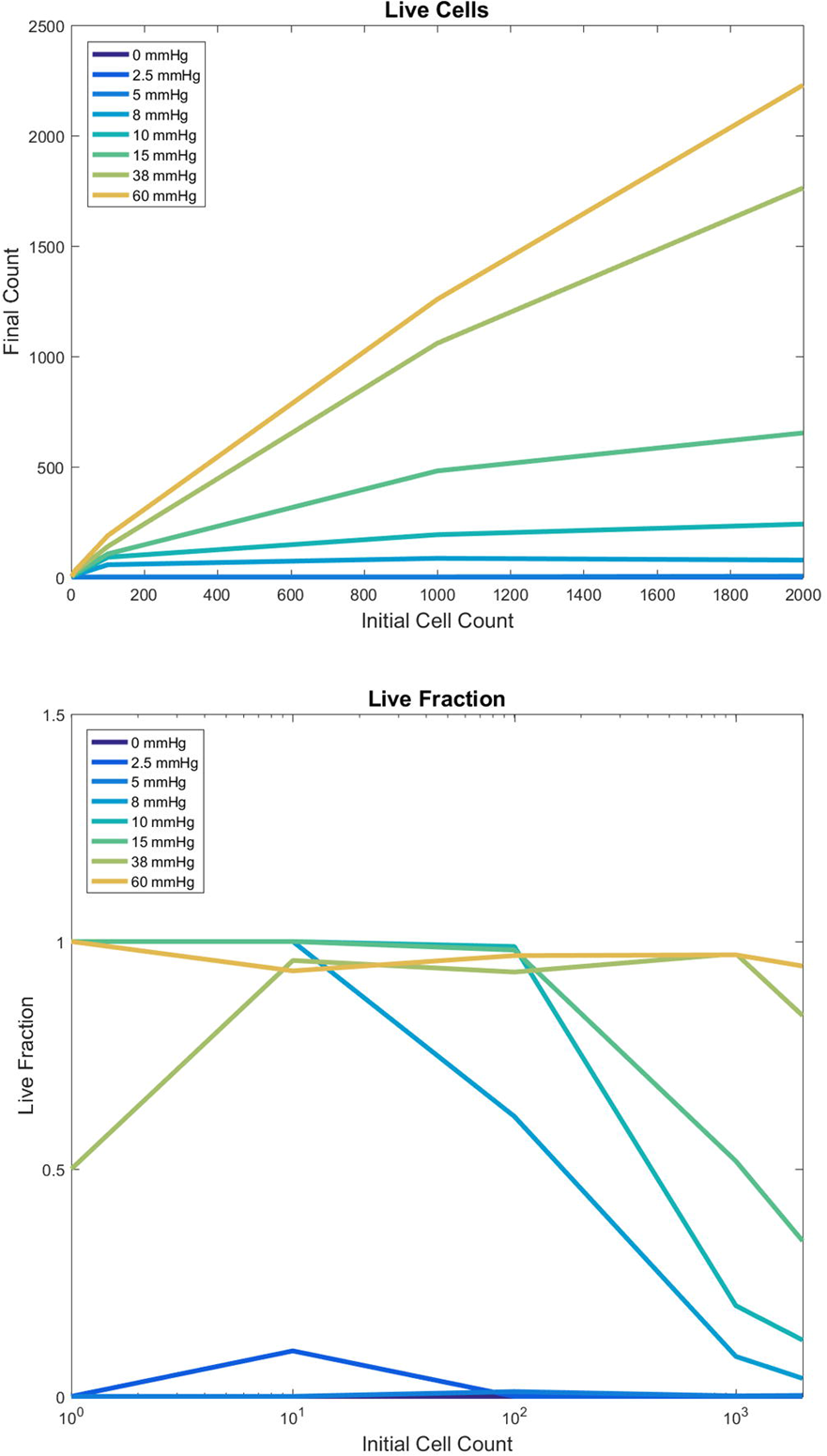
First PhysiCell-EMEWS test on cancer hypoxia: analytics. Live tumor cell count (top) and live cell fraction (bottom) after 48 hours, as a function of oxygenation conditions (each curve is a different condition) and initial cell count (horizontal axis). For intermediate oxygenation conditions, increasing the initial cell count increases the final live cell count (top) but decreases the live cell fraction (bottom). Once oxygenation is high enough, any initial cell count yields nearly 100% live fraction at 48 hours.

### Large-scale cancer immunology investigation

In [32], we performed a single 3-D cancer-immune simulation as detailed above in Sample 3-D simulation. As discussed in [32], the simulation revealed that immune cell homing and tumor-immune interactions are highly non-intuitive, and that immune cell motility parameters play a critical role in the success or failure of the immune response. Had the immune cell “homing” been weaker (i.e., more random, less biased along the chemical gradient), there would have been more mixing between the immune and tumor cells, leading to more cell-cell interactions, a greater probability of tumor cell killing, and a greater effective response. Thus, a broader investigation of the immune cell motility model was warranted.

#### Defining the simulation investigation

We identified the following three model parameters as initial targets for study:

1. **Immune cell attachment rate *rA*:** If an immune cell is in physical contact with a tumor cell, this parameter gives the rate at which they form an adhesive attachment. In any time interval [*t, t* + Δ*t*], the probability of adhering is *rA*Δ*t*. In [32], we set *rA* = 0.2 min*−*1, giving a mean time to attachment of 5 min. <u>Study values:</u> 0.033 min*−*1, 0.2 min*−*^1^, 1.0 min*−*^1^
2. **Immune cell attachment lifetime *TA*:** An attached immune cell that has not successfully triggered tumor cell apoptosis will maintain its attachment for a mean time of *TA*. In any time interval [*t, t* + Δ*t*], the probability of detachment is Δ*t/TA*. In [32], we set *TA* = 60 min. <u>Study values:</u> 15 min, 60 min, 120 min
3. **Migration bias *b*:** Unadhered immune cells choose a motility direction d

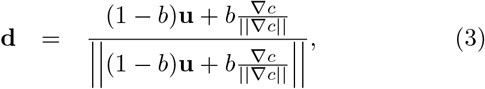

where *c* is the immunostimulatory chemokine and u is a randomly oriented unit vector. Thus, *b* = 0 represents pure Brownian motion, and *b* = 1 represents deterministic chemotaxis along *∇c*; see [32]. We used a default bias *b* = 0.5. <u>Study values:</u> 0.25, 0.50, 0.75

For each of these three parameters, we seek to investigate low, medium and high parameter values, giving a total of 33 parameter combinations. Because the PhysiCell model is stochastic, we seek 5-10 simulations per parameter set, for a total of 135 to 270 simulations. The single sample simulation required approximately 2 days on a four-year-old desktop workstation, including time to save simulation outputs once every three simulated minutes. Thus, our simulation study—performed on a single desktop workstation— would require 270 to 540 days of continuous compute time. Prior to PhysiCell-EMEWS, such a simulation study would be computationally prohibitive.

#### Computational implementation

The parameter sweep implementation was generated using the EMEWS sweep template [52] which allows a user to create an EMEWS project customized for a parameter sweep from the command line. (Additional templates exist for creating ME projects that utilize R or Python ME algorithms.) The project consists of a standard directory structure for organizing model input, output, model launch scripts, and workflow code. The workflow code, implemented in Swift/T, takes as input a text file that explicitly defines all the parameter sets over which to sweep, one parameter set per line. The workflow iterates over each line in the file in parallel and launches a model for each parameter set, taking advantage of the available concurrency. For example, given *n* available processes, *n* models will be run concurrrently. The workflow code can potentially modify the parameter sets, for example, generating additional experimental trials by creating multiple new sets from an existing set through the addition of random seeds. The workflow itself is launched from a bash script which contains place holder values for HPC machine configuration (e.g., queue type, walltime, and so forth), and the parameter input file path. Models and scientific applications such as PhysiCell models are run as Swift/T *app* invocations. An *app* invocation calls out to the external shell to run a bash script that then launches the model. The model launch bash script provided by the EMEWS sweep template takes as arguments the parameter line and a unique directory in which to run the model. The script then runs the model in this directory, passing it the parameter line. It is also possible to run an application as an inmemory Swift/T extension.

For the experiments in this study, the parameter file contained 270 parameter sets. Each parameter set corresponded to a single model running in its own sandboxed directory. The experiments were performed on the Cray XE6 Beagle at the University of Chicago, hosted at Argonne National Laboratory. Beagle has 728 nodes, each with 2 AMD Operton 6300 processors, each having 16 cores, for a total of 32 cores per node; the system thus has 23,296 cores in all. Each node has 64 GB of RAM. Each model was run on a single node, allowing for maximal use of the available threads and the full workflow utilized 272 nodes. 270 were used for model runs, providing complete concurrency while the remaining 2 were used for workflow execution. The workflow completed in 51 hours for a total of 1632 core hours.

#### Simulation results and clinical insights

Using PhysiCell-EMEWS, we initiated 270 simulations of 14 days of growth, followed by a week of immune response: 27 biophysical parameter sets, each with 10 random seeds. Because frequent data saves would significantly slow the simulations due to networked file I/O[32], we only saved the final simulation output for each run, along with SVG visualizations of the *z* = 0 cross-section at intermediate times. Of the 270 requested simulations, 231 were completed in approximately 2 days; see the Discussion for the runs that did not complete.

For each biophysical parameter set, we computed the mean number of live tumor cells remaining at 21 days for the 5-to-10 completed simulation replicates. In Figure 5, we fix the attachment rate at *rA* = 0.2 min*−*^1^ and plot a heat map of this simulation metric versus the migration bias *b* (horizontal axis) and attachment time *TA* (vertical axis)—along with representative tumor cross-sections (i, ii, iii, and iv)—at the final simulation time (21 days). Each shaded square represents the mean live tumor cell count for the *n* simulation replicates (labeled on each square) for a particular parameter set, shaded from deep blue (lowest cell count; most effective response) to bright yellow (highest cell count; least effective response).

**Figure 5.**
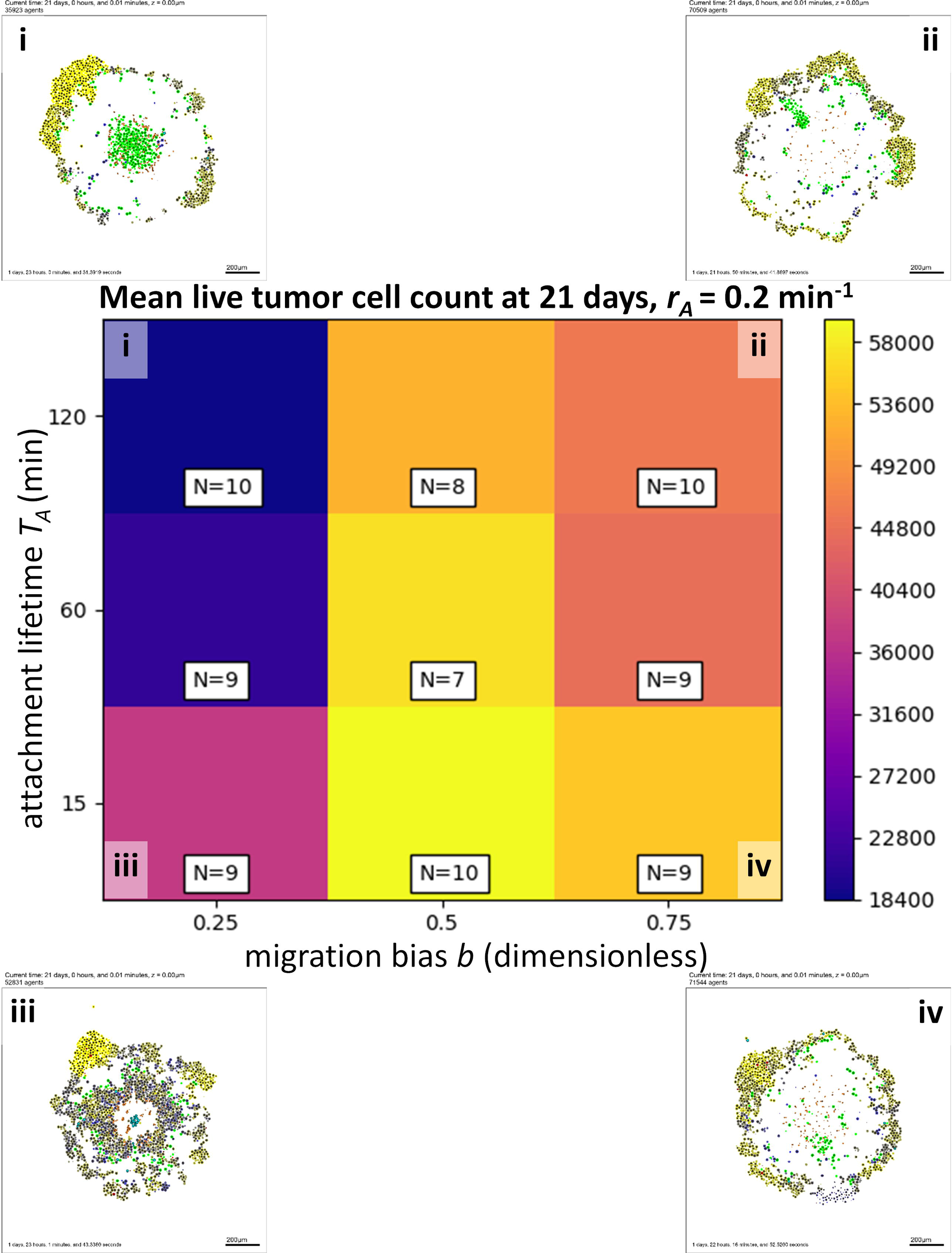
High-throughput 3-D cancer-immune simulation: impact of migration bias and and attachment lifetime. We plot a heatmap for final live cell tumor count (blue is lowest, or most effective immune response; yellow is worst immune response) for varied migration bias (horizontal axis) and immune cell attachment lifetime (vertical axis). Characteristic final tumor cross sections are labeled i-iv. In particular, decreasing migration bias improves the response.

For all values of *TA*, decreasing the migration bias (and thus decreasing homing along the immunostimulatory gradient) dramatically improved the immune response. This result was slightly non-intuitive, as it suggests that the efficiency and precision of chemotaxis, if maximized, leads to an “overshoot” phenomenon that actually works against the goal of increasing tumor-immune cell mixing, an important factor in the ability to kill tumor cells noted in [32]. Alternatively, for any fixed migration bias *b*, increasing the attachment lifetime also improved the immune response as would be expected, although increases beyond 60 minutes were only marginally helpful. However, these results demonstrate the need to account for different axes of affect in any attempt to optimize towards a particular goal (e.g., a therapeutic design goal of maximizing tumor-immune cell mixing to increase tumor cell killing).

In Figure 6, we show a heat map for the mean live tumor cell count at 21 days versus migration bias *b* (horizontal axis) and the attachment rate *rA* (vertical axis). For all values of *b*, increasing the attachment rate improved the response, although the improvement beyond 0.2 min*−*1 was marginal. Interestingly, for a fixed attachment rate *rA*, the impact of *b* was non-monotonic. Either decreasing *b* (to promote random tumor-immune mixing) or increasing *b* (to allow more directed cell migration) would improve the immune response over the initial value of 0.5. This again highlights the nonlinear nature of tumor-immune interactions, and the need for high-throughput investigation of mechanistic 3-D models to systematically probe these dynamics and identify trade-offs that need to be accounted for when designing putative therapies.

**Figure 6.**
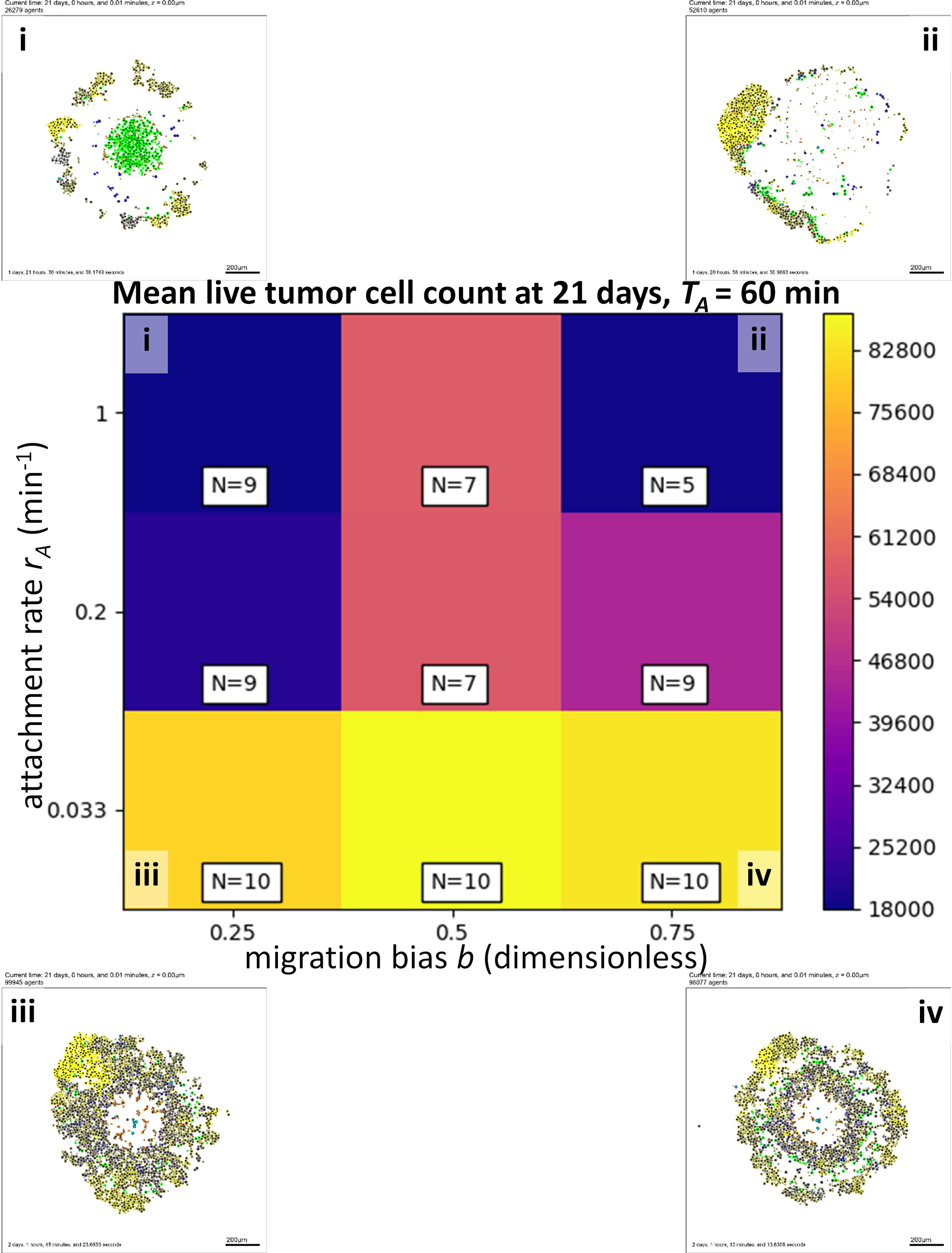
High-throughput 3-D cancer-immune simulation: impact of migration bias and and attachment rate. We plot a heatmap for final live cell tumor count (blue is lowest, or most effective immune response; yellow is worst immune response) for varied migration bias (horizontal axis) and immune cell attachment rate (vertical axis). Characteristic final tumor cross sections are labeled i-iv. The impact of both parameters was nonlinear.

In Figure 7, we show a heat map for the mean live tumor cell count at 21 days versus the attachment rate *rA* (horizontal axis) and the attachment lifetime *TA* (vertical axis), with *b* = 0.5. For all attachment lifetimes *TA*, increasing the attachment rate improved the immune response, as expected. However, for higher attachment rates *rA*, there was an interesting trend towards bimodal optima when examining the impact of the attachment lifetime: increasing the attachment lifetime from the medium (1 hour) to high (2 hour) value improved the treatment response, possibly by increasing the likelihood of a successful apoptosis event for any tumor-immune cell-cell attachment. However, decreasing the attachment lifetime from medium (1 hour) to short (15 minutes) *also* improved the response, likely by increasing the number of tumor-immune cell-cell attachments. This demonstrates that the highly nonlinear dynamics of the cancer-immune interactions can admit many potential therapeutic strategies, some of which may be non-intuitive. Additional simulations are planned to determine whether this is an artifact of low replicate numbers, or represents an actual non-normal distribution in the dynamic range of these parameters.

**Figure 7.**
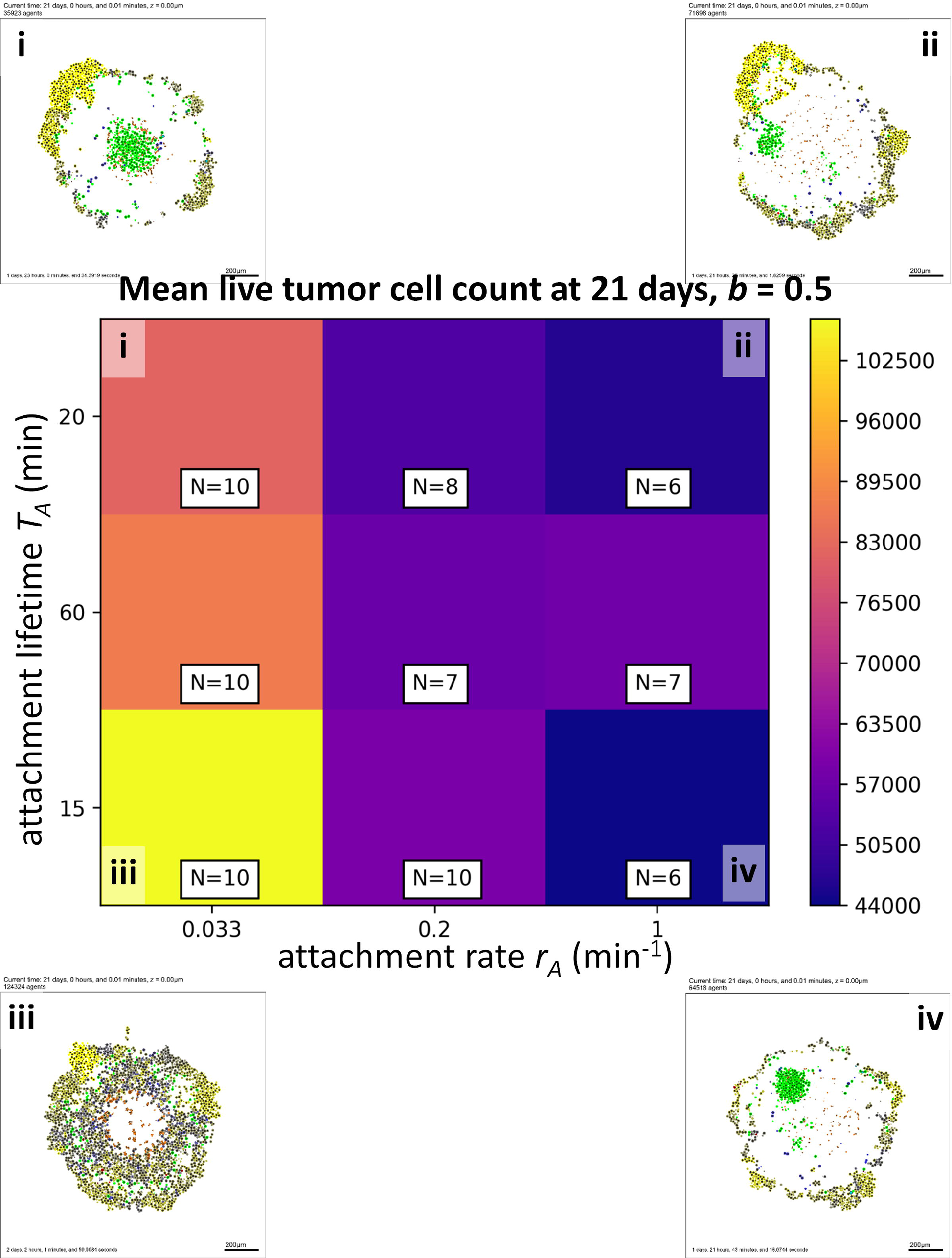
High-throughput 3-D cancer-immune simulation: impact of attachment rate and and attachment lifetime. We plot a heatmap for final live cell tumor count (blue is lowest, or most effective immune response; yellow is worst immune response) for varied immune cell attachment rate (horizontal axis) and immune cell attachment lifetime (vertical axis). Characteristic final tumor cross sections are labeled i-iv. The impact of both parameters was nonlinear.

## Discussion

Despite the prototyping nature of these simulation experiments, we believe that there are important insights that can be gained by these results. Most significant is substantiation of the general belief that multidimensional, nonlinear systems can lead to some nonintuitive results. In the context of cancer immunology, we found that reducing chemotactic efficiency (reducing attraction bias) can actually be beneficial in terms of achieving an intermediate goal (tumor-immune cell mixing) that improves the functional output (tumor suppression). Additionally, these results, while qualitative in nature, suggest that many immunotherapy design parameters have thresholds values, beyond which further refinements give little or no clinical benefit.

The identification of seeming thresholds for therapeutic parameters such as attachment duration and rate suggests that higher resolution models may be used to identify boundary conditions for future wet lab experimental investigations, which in turn can be used to refine the computational models in exactly the type of iterative workflow envisioned in Figure 2. At some point, the results from this workflow will aid in “pre-screening” potential research spending priorities away from target goals where further improvements (i.e., to speed up the attachment rate or increase the attachment lifetime) *would not* improve the immune response. In cases of non-monotonic system behavior (e.g., where either high or low migration bias can lead to treatment success, whereas intermediate migration bias yields a poorer outcome), high-throughput model investigations may be all the more critical to identifying robust treatment designs with more reliable patient outcome.

While the current model yielded fresh insights on cancer-immune interactions, further refinements are needed to unlock its full potential. In future work, we plan to integrate and explore other key features of the immune system, such as inflammatory responses, cross-talk between different immune cell types, and molecular-level mechanisms for MHC function and immune-mediated cancer cell apoptosis [53, 47, 48, 21]. The models also need extension to directly model new treatments such as the role of PD-1 and PD-L1 in CAR T-cell therapies [19, 20]. In our next steps, we will extend the modeling framework to incorporate these effects, and import it into the EMEWS framework. We will start exploring the emergent tumor response to immune therapy under a variety of immune cell hypotheses and cancer phenotypes. Ultimately, we will generate hypotheses that elucidate the most and least ideal patient characteristics for immunotherapies.

In our pilot work to date, we have run a *single iteration* of the hypothesis testing loop; our next step is to complete the loop and iteratively optimize the treatment response over the current “design” parameters (attachment lifetime, migration bias, and attachment rate). This should yield testable hypotheses on immune system conditions for effective and ineffective tumor suppression. We also plan separate cancer hypothesis investigations in the PhysiCell-EMEWS framework. In ongoing breast cancer projects, we are evaluating families of cell-cell interaction hypotheses for “leader cells” (highly motile, less proliferative) and “follower cells” (less motile, more proliferative) that best explain time series morphologic data [54]. This work will further test the potential of PhysiCell-EMEWS to not merely explore large parameter spaces, but to optimally match hypotheses to experimental observations. We would then develop independent experiments to validate or refine the optimal hypotheses.

We note that the generalized description of hypotheses is not yet mature. Standards have emerged to describe molecular-scale systems biology (generally systems of ODEs) as SBML [55], and more recently to express multicellular biology as MultiCellDS, but cell-cell interaction rules will likely require a different description, such as by using elements of the Cell Behavior Ontology [56].

PhysiCell-EMEW’s computational performance could be further improved. In particular, the diffusion solver (BioFVM) is well-suited to leveraging GPU resources available on today’s typical HPC/HTC compute nodes using, for example, OpenACC, CUDA, or OpenCL. Scientifically, complex molecular-scale systems biology is typically written as SBML (systems biology markup language) models, and so to integrate these into high throughput multiscale mechanistic hypothesis testing, we plan to implement an SBML model integrator, such as the cross-platform libRoadrunner platform [57].

Lastly, we note that there were other benefits to combining PhysiCell and EMEWS to run a large number of simulations: we estimate that the cancer-immune investigation included on the order of 1 to 100 billion calls to the of the tumor-immune mechanical and biochemical interaction codes. This allowed us to “stress test” PhysiCell and identify rare bugs for future code releases. 39 simulations in our investigation terminated prematurely due to rare events, such as multiple immune cells attempting to apoptose the same tumor cell, or a tumor cell necrosing while still attached to an immune cell; these rare events removed dead cells from memory while memory pointers were still in active use, occasionally causing segmentation faults. Without high-throughput simulation investigations (which included over a year of compute time), these bugs would likely remain undetected and unfixed for years. We anticipate that other open source computational biology projects could similarly benefit from high-throughput testing in EMEWS.

## Conclusions

We have demonstrated a 3-D mechanistic tumor-immune interaction model (and more generally, a mechanistic agent-based cancer modeling platform, using PhysiCell) that has an appropriate balance of flexibility, efficiency, and realism for efficient single simulations, that predict the emergent systems behaviors for a given set of cancer hypotheses. It is self-contained code (can be distributed as a ZIP file) enabling very simple deployment.

We have shown how a previously-developed extreme-scale model exploration and optimization platform (EMEWS) can compatibly deploy PhysiCell for model exploration in high throughput. We have outlined the overall platform to perform high-throughput hypothesis testing on using PhysiCell and EMEWS, and we gave an early example on a simple (but spatially nontrivial) model system of hypoxic tumor growth. We then demonstrated PhysiCell-EMEWS with a large parameter space investigation of a mechanistic 3-D cancer-immune model, obtaining significant and nonintuitive insights on immune cell homing and adhesion dynamics that would not have been feasible without HTC. The next natural step is to iterate past this first investigation and find therapeutic design optima that maximize tumor regression; this would represent a full test of PhysiCell-EMEWS as a hypothesis optimization tool.

Cancer biology—particularly cancer-immune interactions—occurs in complex dynamical, multiscale systems that frequently yield surprising emergent behaviors that can impair treatment. High-throughput model investigation and hypothesis testing affords a new paradigm to attacking these complex problems, gaining new insights, and improving cancer treatment strategies.

We close by noting that this framework has applications beyond cancer. In general, testing multiscale hypotheses in high throughput is valuable in determining the rules underlying (often puzzling) experimental data, and even to evaluate the limitations of experiments themselves [29, 30]. The PhysiCell-EMEWS system could be used as a multicellular design tool: for any given multicellular design including single-cell and cell-cell interaction rules (which map onto hypotheses in this framework), PhysiCell-EMEWS can test the emergent multicellular behavior against the target behavior (the design goal), and iteratively tune the cell rules to achieve the design goal. In [32], we began to design cell-cell interaction rules to create a multicellular cargo delivery system to actively deliver a cancer therapeutic beyond regular drug transport limits to hypoxic cancer regions. In that work, we manually tuned the model rules to achieve this (as yet unoptimized) design objective, requiring weeks of people-hours to configure, code, test, visualize, and evaluate. Integrating such problems into a high-throughput design testing system such as PhysiCell-EMEWS would be of clear benefit.

## Abbreviations

ABM (agent-based model): A computational model focused on independent (but often interacting) software agents;
BLAS (basic linear algebra subroutine): fundamental linear operators, such as linear addition of vectors;
CAR (chimeric antigen receptor): a type of engineered receptor (usually on T cells) binding a tailored specificity to an effector immune cell;
EMEWS (Extreme-scale Model Exploration with Swift): a framework for model exploration using the Swift/T parallel scripting language;
HPC (high performance computing): solution of large and complex problems by parallelization over networked computers, generally supercomputers;
HTC (high throughput computing): the use of many computing resources over long periods of time (not necessarily linked to a high-speed network as in HPC).
LOD (locally one-dimensional): a method for numerically solving partial differential equations based upon solving lower-dimensional problems;
ME (model exploration): combinatorial mixing of a model’s parameter values;
MHC (major histocompatibility complex): a cell surface molecule used by immune cells to identify foreign cells;
ODE (ordinary differential equation): an ordinary differential equation;
PDE (partial differential equation): a partial differential equation;

## Author’s contributions

GA and PM initiated and designed the overall PhysiCell-EMEWS project. SHF, AG, and PM designed and developed the original PhysiCell software. PM developed the cancer-immune model. JO, NC, JMW, CM, CC and GA designed, developed and executed the EMEWS framework for this project. RH and PM analyzed the resulting data and provided the figures. GA and PM obtained funding. All authors contributed to, read, and approved the final version of this manuscript.

## Ethics approval and consent to participate

Not applicable.

## Consent for publication

Not applicable.

## Availability of data and materials

All simulation source code and scripts for execution and analysis for this project (including data generation) are available at https://github.com/MathCancer/PhysiCell-EMEWS and at [58].

